# Seo1p, a high affinity, plasma membrane transporter of the γ-Glu-met dipeptide in yeasts and fungi

**DOI:** 10.1101/2025.01.15.633159

**Authors:** Pratiksha Dubey, Md Shabbir Ahmad, Sunil Laxman, Anand K Bachhawat

**Affiliations:** Department of Biological Sciences, Indian Institute of Science Education and Research, Mohali, SAS Nagar, Punjab 140306 India; DBT- Institute for Stem Cell Science and Regenerative Medicine (inStem), Bangalore, India

## Abstract

γ-Glu dipeptides are ubiquitous in nature, and yet their metabolism and transport are poorly understood. Here we investigate this using the dipeptide γ-Glu-met in *Saccharomyces cerevisiae*. γ-Glu-met was efficiently utilized by *S. cerevisiae* and its degradation was dependent on both the glutathione degrading cytosolic Dug2p/Dug3p complex, and the vacuolar Ecm38p. Using a transcriptomics approach, followed by a genetic screen, we identified Seo1p, an orphan transporter of yeast, as the transporter of γ-Glu-met. Uptake studies confirmed Seo1p as a high affinity (Km =48uM), highly specific transporter of γ-Glu-met since other analogs like n-Glu-met, γ-Glu-leu, γ-Glu-cys, γ-Glu-met-gly, methionine and methionine sulfoxide were not transported by Seo1p. Candida spp. also encoded a functional Seo1p. A second transporter, Opt2p, identified in the screen, was also investigated. However, Opt2p was not primarily involved in γ-Glu-met uptake. Its deletion affected vacuolar morphology, that interfered with the degradation of the peptide through Ecm38p. These studies demonstrate how organisms have evolved dedicated pathways for the uptake of these unusual peptides.

## Introduction

γ-Glutamyl peptides are a class of widely prevalent, low-molecular weight peptides of 2−3 amino acids, where an N-terminal glutamic acid links to an amino acid or a peptide. Some of the γ-Glutamyl tri-peptides are well known and extensively investigated. For example, the γ- Glutamyl tri-peptide, glutathione (γ-Glu-cys-gly) that is critical for redox and other functions is very abundant in eukaryotes. The tri-peptide homoglutathiones that include γ-Glu-cys-ser and γ-Glu-cys-β-ala have been found in considerable amounts in certain plants (Klapheck, 1988), and the glutathione analogue, ophthalmic acid (γ-Glu-2-aminobutryl glycine) has been known for many years (Schomakers et al., 2024). Much less is known about the γ-Glutamyl di- peptides. Of these, the γ-Glutamyl di-peptide, γ-Glu-cys, which is a precursor of glutathione and is the thiol-equivalent of glutathione in many archaea is the best studied (Newton and Javor, 1985). However, there are several γ-Glu-di-peptides that are ubiquitously present, across the kingdoms of life, including mammals, plants, fungi, yeast and bacteria. One study identified many different γ-Glu peptides that included γ-Glu-glu, γ-Glu-met, γ-Glu-phe, γ-Glu-tyr, γ-Glu-trp and γ-Glu-leu in cyanobacteria (Jaiswal et al., 2022). Indeed, γ-Glu-di-peptides have been identified in many different organisms including mammals (Cassier-Chauvat et al., 2023; Iciek et al., 2009; Sofyanovich et al., 2019; Thacker et al., 2021).

The γ-Glutamyl-di-peptides are thought to be synthesized through enzymatic side reactions of different enzymes that includes γ-Glutamyl transpeptidase (GGT), γ-Glutamyl cysteine synthetase or γ-Glutamyl cysteinyl ligase (GCS or GCL), glutathione synthetase (GS) and glutaminase (Klapheck, 1988; Soga et al., 2011; Suzuki et al., 2007). In case of γ-Glutamyl cysteinyl ligase, limiting levels of cysteine (which is often observed in cells) results in alternate amino acids being accepted by the enzyme that leads to various γ-Glutamyl-dipeptide formations (Kelly et al., 2002). As a consequence of being side reactions of various enzymes, the γ-Glutamyl-di-peptides have been considered metabolic byproducts of different enzymes, and their physiological significance was for many years unclear. However, recent evidences have suggested that γ-Glu-di-peptides have physiological roles in cells. Many γ-Glu-peptides impart kokumi taste to food by activating the extracellular calcium-sensing receptor (CaSR) (Ohsu et al., 2010). The γ-Glutamyl dipeptide γ-Glutamyl glutamate has an excitatory effect on synaptic transmission by activating N-methyl-D-aspartate receptors in human and rat (Sebih et al., 2021). Other γ-GPs have anti-inflammatory, antioxidant, metal ion chelating, antitumor, and hypoglycemic effects (Guha and Majumder, 2022), and some γ-Glutamyl-dipeptides may have therapeutic importance (Yang et al., 2018b) and disease associations (Soga et al., 2011; Xiao et al., 2017; Zierer et al., 2016).

The γ-Glutamyl-dipeptides are very stable due to the γ-Glutamyl bond, making them resistant to peptidases, and their degradation requires specific enzymes. Mammalian cells and bacteria contain γ-Glutamyl-cyclotransferases (γ-GCT) which breaks down a γ-Glu-amino acid into 5-oxoproline and the amino acid. However, these enzymes are absent in fungi and plants, so their metabolism in these organisms is not known. How these peptides traverse across the cell membrane and enter the cell are also not known in different organisms. In humans, multiple pathways have been postulated for the transport of these peptides across the intestinal epithelial cell monolayer. These include active transport mediated by the peptide transporter 1 (PepT1), the paracellular route through tight junctions, transcytosis, and passive transcellular diffusion (Xu et al., 2019). PepT1 can facilitate the transportation of a variety of small peptides, such as Gly-Sar, Trp-His, Gly-Pro, Ile-Arg-Trp (Bejjani and Wu, 2013). However, while the transport of γ-Glu-Val as well as γ-Glu-S-(Me) C, γ-Glu-Leu via PepT1 have been suggested, γ-Glu-di- peptide transport by specific pathways has not yet been demonstrated.

Given this large gap in knowledge, we have initiated a study on γ-Glu-di-peptides in yeast to investigate the transport and metabolism of a γ-Glu-di-peptide in *S. cerevisiae*. To address this question, we chose γ-Glu-met as the peptide, for two reasons. First, it has a physiological role in cells (Yang et al., 2018a). Second, it enabled us to set up a simple genetic screen to facilitate the analysis. We exploited the *S. cerevisiae met15Δ* strain (an organic sulfur auxotroph), where we observed that γ-Glu-met could be used as a source of sulfur after its transport and cleavage to methionine. The known peptide transporter, Ptr2p and the known glutathione /oligopeptide transporter, Hgt1p/Opt1p, did not have a role in its transport. Hence, we attempted to identify both the transporter of γ-Glu-met, and the enzymes involved in its degradation. Our studies reveal that the degradation of γ-Glu-met occurs through the action of two glutathione degrading enzymes, Dug2p/Dug3p in the cytoplasm and the Ecm38p γ-glutamyl transpeptidase in the vacuole. We further identify an orphan transporter of *S. cerevisiae*, Seo1p as an evolutionarily conserved, specific high affinity transporter of γ-Glu-met. This study substantially advances our understanding of how these ubiquitous and unusual dipeptides might be transported, and metabolized.

## Results

### Yeast cells can efficiently utilize γ-Glu-met as a sulfur source through degradation by the Dug2p/Dug3p protein complex and Ecm38p

To investigate the utilization of γ-Glu-met in yeast, we exploited the *met15Δ* strain, an organic sulfur auxotroph. The *met15Δ* strain exhibits growth only upon supplementation with an organic sulfur source such as methionine, cysteine, or glutathione. γ-Glu-met contains methionine, which can serve as a sulfur source. However, for this peptide to act as a sulfur source, it needs to be transported into the cell and subsequently degraded into glutamate and methionine. To test this possibility, we provided γ-Glu-met to *met15Δ* (sulfur auxotroph) cells at a concentration of 100 μM. We observed that *met15Δ* cells could efficiently utilize γ-Glu-met as a sulfur source (Fig1a). The generation time on methionine was 2.0 hrs whereas on γ- Glu-met it was 4.0 hrs. This observation suggests the presence of a transport system for this peptide, as well as a degradation mechanism within the cells.

**Figure 1:**
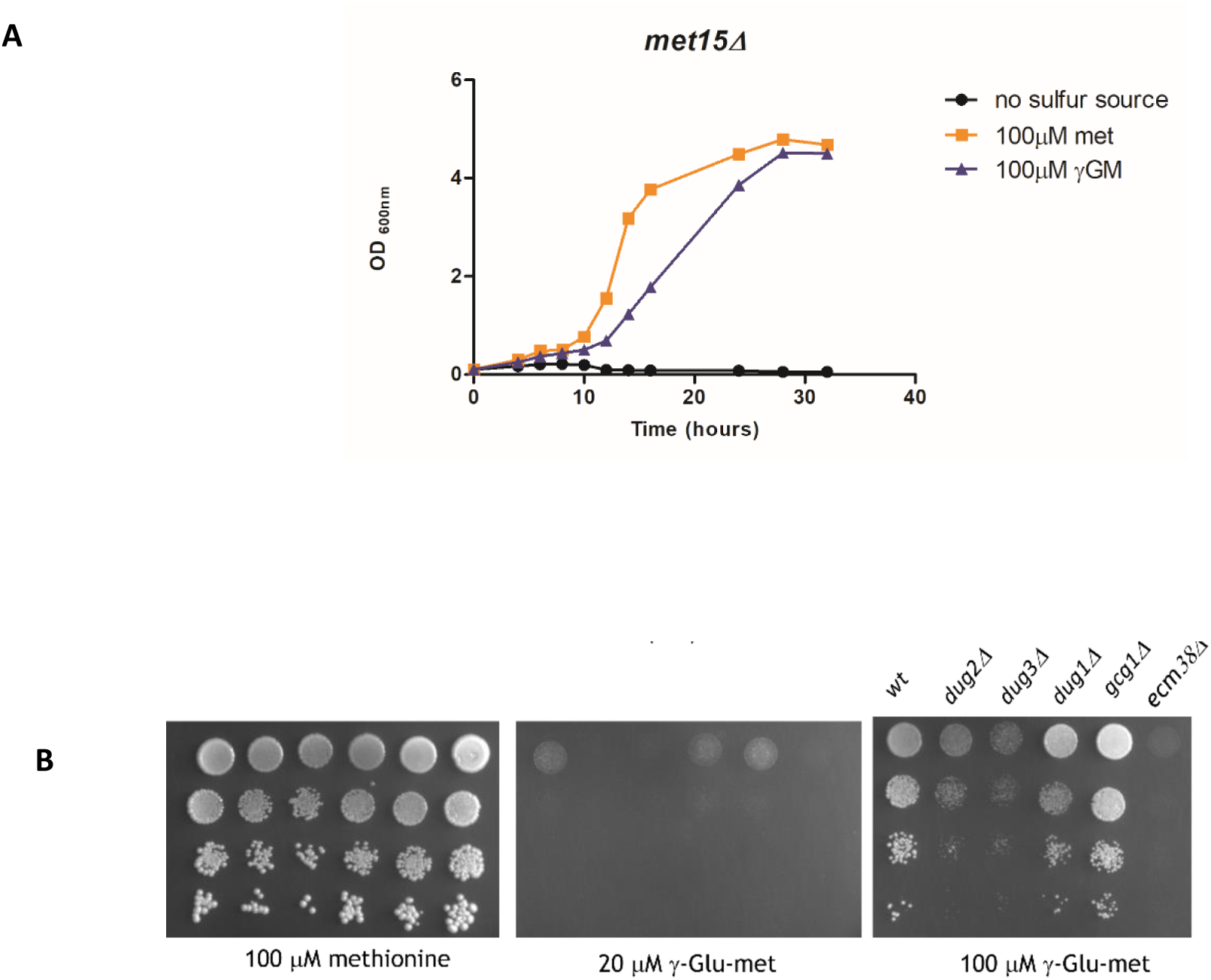
γ-Glu-met as a sulfur source in yeast cells (BY4741) and its breakdown by different enzymes: A) γ-Glu-met and methionine at 100μM conc or no added organic sulfur in SD medium, B) Evaluation of *ecm38Δ*, *dug1Δ, dug2Δ* and *dug3Δ* on γ-Glu-met (100μM); Serial dilutions were spotted on the plates as described in materials and methods.

To find out the pathways involved, we initially evaluated previously known glutathione degrading enzymes since glutathione is a γ-Glutamyl tri-peptide, and previously known transporters for glutathione and peptides. There are four enzymes known to be involved in glutathione degradation, these include the Dug1 peptidase (Kaur et al., 2009) that cleaves the cys-gly peptide and the Dug2-Dug3 complex (Kaur et al., 2012), Ecm38p (γ-Glutamyl transpeptidase) (Kumar et al., 2003) and Gcg1p (Yeast Chac2) (Kaur et al., 2017; Kumar et al., 2012), three enzymes capable of cleaving the γ-Glutamyl bond. The Dug2-Dug3 complex is encoded by two different proteins, Dug2p and Dug3p which combine to make a functional protein complex. The effect of deleting these different genes in a *met15Δ* background were evaluated. All these deletion strains were tested for γ-Glu-met utilization, based on assessing the eventual growth of cells. We find that *dug2Δ*, *dug3Δ* and *ecm38Δ* were defective in utilizing the peptide, and *ecm38Δ* was most severely affected in the utilization as seen on plates (Fig1b) and on liquid medium (Fig S1). This data suggests that these enzymes participate in the degradation of γ-Glu-met in glutamate and methionine. Dug2p-Dug3p is a cytoplasmic complex previously known to be involved in GSH degradation and the finding here suggest that the activity is not restricted to the tripeptide but it can also cleave γ-Glu-dipeptides. Ecm38p on the other hand is present on the vacoular membrane, with its active site facing the vacuolar lumen.

### The putative membrane transporter, Seo1p, is involved in γ-Glu-met transport

In the yeast *S. cerevisiae*, peptide or oligopeptide uptake is driven by different transporters. PTR2 is currently the sole known di and tri peptide transporter in *S. cerevisiae* (Perry et al., 1994), while HGT1/OPT1 is a glutathione/oligopeptide transporter (Bourbouloux et al., 2000; Hauser et al., 2000). When we evaluated the effect of deletion of these genes in the *met15Δ* background, we did not observe any defect in growth (Fig 2a). This suggests that these two transporters are not involved in the utilization of γ-Glu-met.

**Figure 2:**
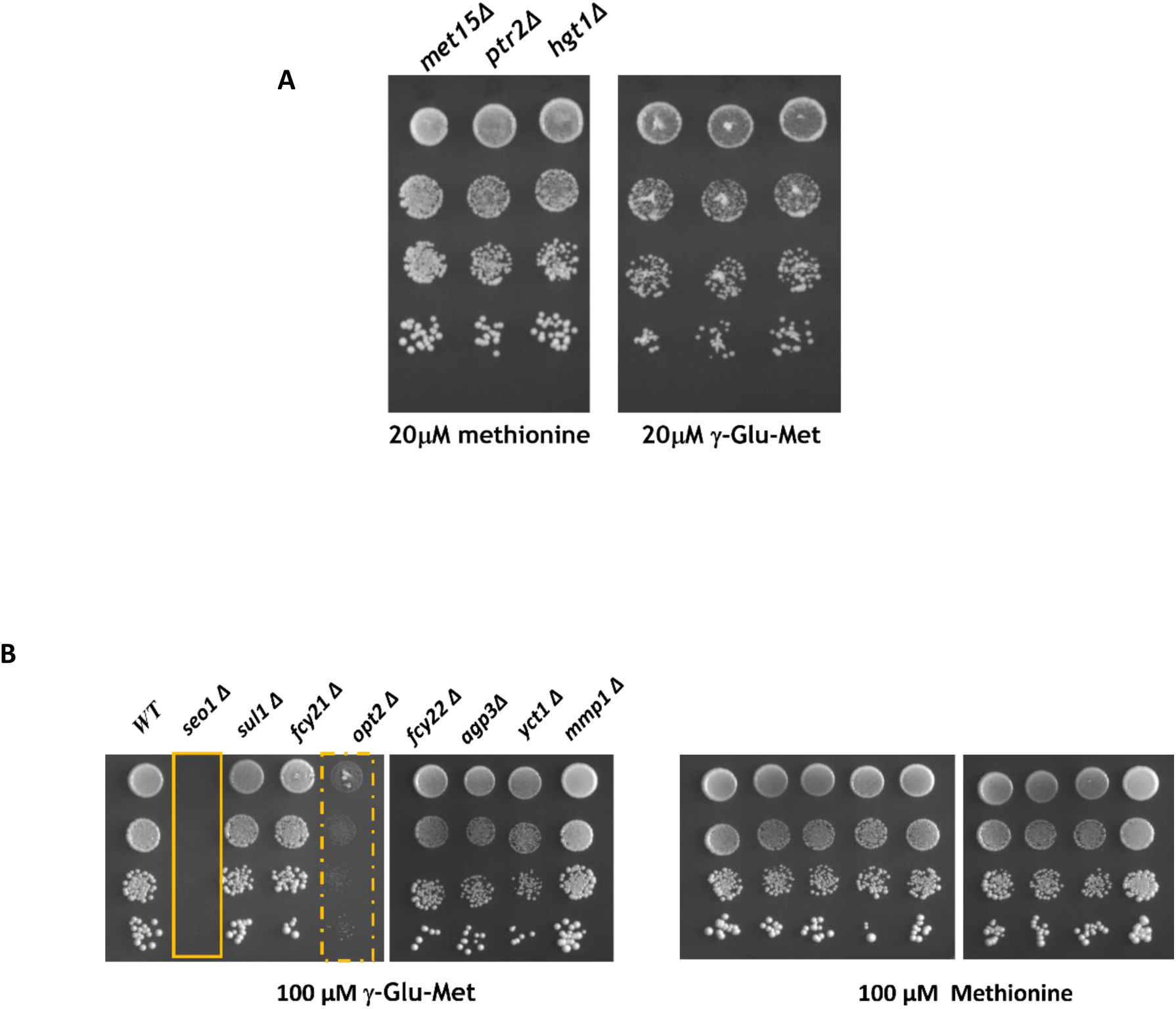
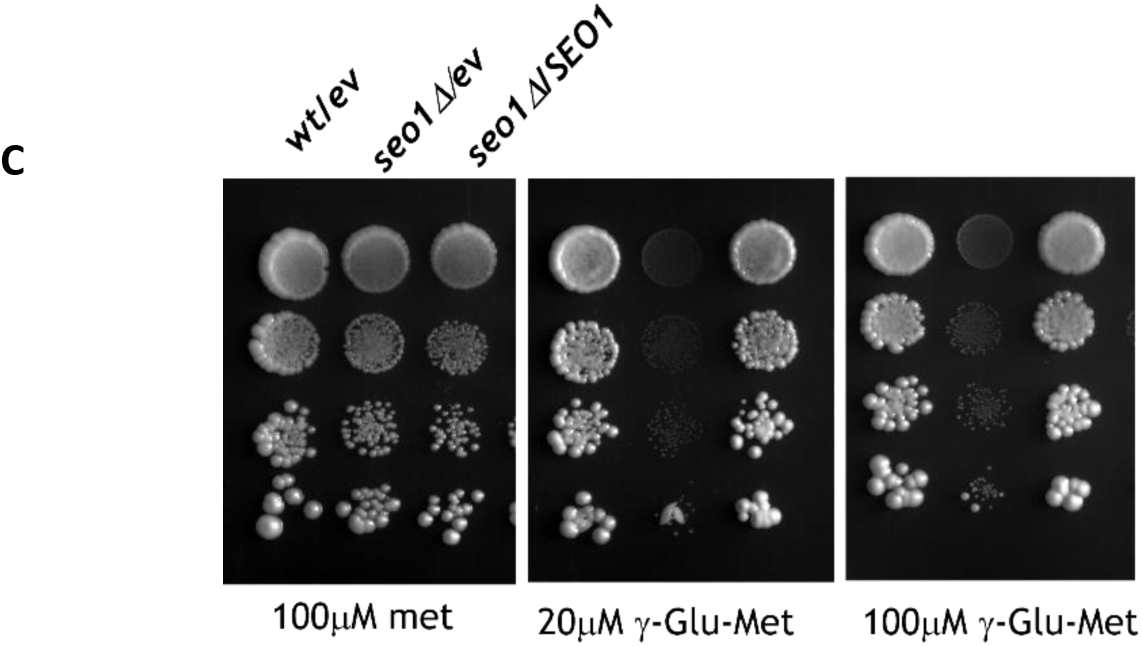
Evaluation of different transporter deletion strains in a *met15Δ* background to utilize γ-Glu-met: A) Growth of deletions of the peptide transporter (*ptr2Δ*) and glutathione transporter (*opt1Δ /hgt1Δ)* in a *met15Δ* background on γ-Glu-met. **B)** The top upregulated transporter deletion strains in *met15Δ* background were assessed for γ-Glu-met utilization **C)** The defective phenotype of *seo1Δ* strain is rescued upon expression of SEO1

As none of the known transporters appeared to be involved in γ-Glu-met transport, we carried out transcriptomic analysis of *S. cerevisiae met15Δ* cells grown on γ-Glu-met as compared to methionine to identify if there were any putative transporters that are upregulated in presence of γ-Glu-met. Cells were grown to mid log phase, harvested and RNA isolated and RNA seq analysis done as described in materials and methods. The total read count for the replicate 1 and replicate 2 of methionine treated cells (control) was 15 million and 12 million respectively, whereas in case of γ-Glu-met treated cells (test), the total read counts for replicate 1 and replicate 2 was 13 million and 16 million respectively. Linear regression analysis done for the deferentially expressed genes displayed a correlation coefficient value r=1.00 and r=0.99 for control and test respectively showing high reproducibility (Fig S3). We observed that the sulfur regulated genes were induced in γ-Glu-met treated cells (Table S1), consistent with an expectation, as methionine is a repressing sulfur source related to γ-Glu-met. We focused our attention, however on the transporter genes (Table 1). Many of the upregulated transporters were also known sulfur regulated proteins (Boer et al., 2003), and we asked if these might be involved in γ -Glu-met utilization, by evaluating the growth on on γ-Glu-met for deletions in a *met15Δ* background .

**Table 1:** Transporter genes upregulated in γ-Glu-met versus methionine as seen by RNA seq. *met15Δ* cells were grown on either γ-Glu-met or methionine as a sulfur source and harvested at 1 OD. RNA seq analysis was carried out to find out the upregulated transporters as described in the Methods. Only the genes which had a P<0.05 and a log2 foldchange >1 were included in the list.

| *protein* | *log2Fold* | *P Value* | *function* |
| --- | --- | --- | --- |
| *SUL1* | 3.75 | 6.67E-35 | High affinity sulfate permease of the SulP anion transporter family |
| *SEO1* | 3.26 | 5.54E-29 | Putative permease |
| *YCT1* | 2.89 | 2.87E-23 | High-affinity cysteine-specific transporter |
| *AGP3* | 2.50 | 1.83E-18 | Low-affinity amino acid permease |
| *SUL2* | 2.25 | 5.45E-16 | High affinity sulfate permease |
| *MEP2* | 1.45 | 4.83E-07 | Ammonium permease involved in regulation of pseudohyphal growth |
| *VBA5* | 1.31 | 0.022912 | Plasma membrane protein of the Major Facilitator Superfamily (MFS) |
| *ATR2* | 1.22 | 0.00065 | Putative boron transporter involved in boron efflux and resistance |
| *FCY21* | 1.11 | 0.000123 | Putative purine-cytosine permease |
| *OPT2* | 1.04 | 0.000219 | Oligopeptide transporter |
| *MMP1* | 1.03 | 8.97E-05 | High-affinity S-methylmethionine permease |

We observed that *seo1Δ* showed a complete growth defect on γ-Glu-met supplemented synthetic medium, while *opt2Δ* also showed a fairly severe growth defect (Fig 2b). As Seo1p is an uncharacterized transporter, we initially focused on SEO1. The *seo1 Δ* phenotype could be rescued by complementation with a WT copy of SEO1 (Fig 2c).

To confirm if γ-Glu-Met uptake required Seo1p we carried out uptake studies, where intracellular γ-Glu-Met levels were measured by LC-MS/MS. For this study the *seo1Δ met15Δ ecm38Δ dug3-2* strain, that is deficient in γ-Glu-Met degradation (as described above), was used. *SEO1* (under a TEF promoter) was expressed in these strains. The SEO1 overexpression in *seo1Δ met15Δ ecm38Δ dug3-2* strain showed a time dependent uptake of γ-Glu-Met (100uM) (Fig 3a), while negligible uptake was consistently observed with the control vector. These data unambiguously reveal that SEO1 is involved in γ-Glu-Met transport. Next, we determined the effective Km for γ-Glu-Met transport, by measuring uptake at fixed times, for different concentrations of γ-Glu-Met, and found that an effective Km was 48μM (Fig 3b). This was very consistent with the rescue of growth seen with growth of cells in ∼100 uM γ-Glu-Met.

**Figure 3:**
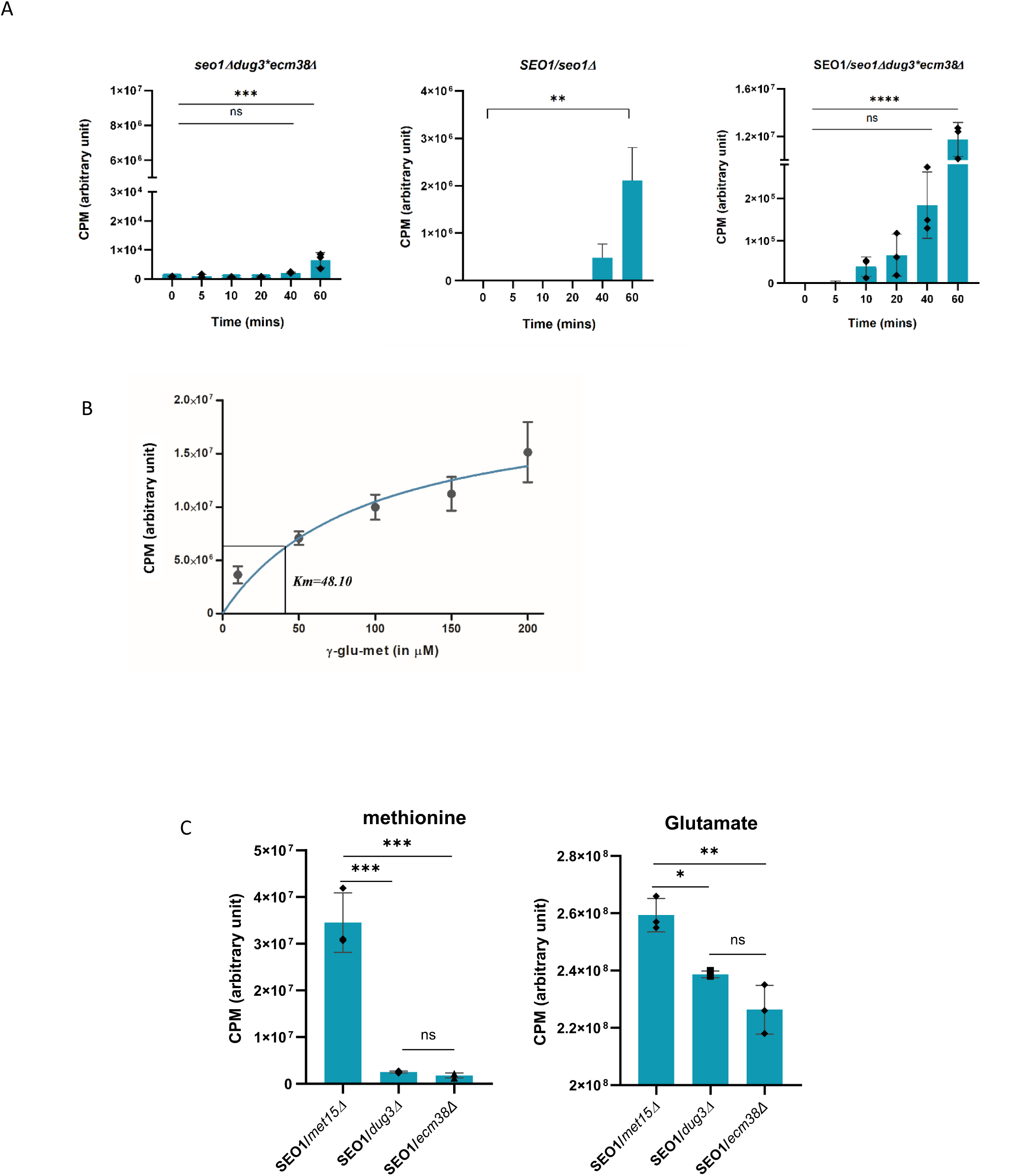
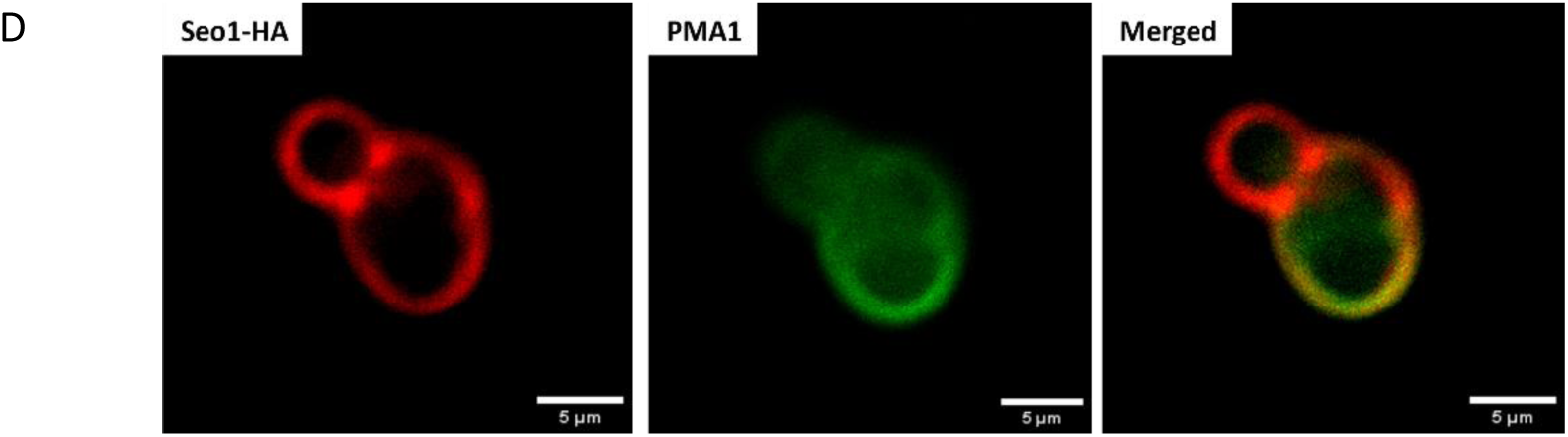
Seo1p is a plasma membrane transporter of γ-Glu-Met in yeast A) γ-Glu-Met uptake was measured using LC-MS/MS at different time intervals i.e. 5, 10, 20,40 and 60 min. left panel shows the vector/*seo1Δmet15Δecm38Δdug3-2* strain with no γ-Glu-Met uptake. SEO1 overexpression in *seo1Δmet15Δecm38Δdug3-2* shows gradual increase in γ-Glu-Met levels, maximum uptake was seen at 60mins. **B)** Different concentrations of γ-Glu-Met i.e. 10, 50, 100, 150 and 200 μM was used for uptake by SEO1 in *seo1Δmet15Δecm38Δdug3-2* Strain and Km was determined by michelis-menton equation (graph pad Prisma). **C)** γ-Glu-Met degradation profile was assessed by methionine and glutamate levels in *met15Δ, ecm38Δ and dug3Δ,* lower levels of methionine in *ecm38Δ and dug3Δ* was found in comparison to *met15Δ* due to the defective degradation of γ-Glu-Met into methionine. Glutamate levels were also reduced in *ecm38Δ and dug3Δ* cells in comparison to *met15Δ.* **D)** cellular localization of SEO1: anti-HA antibody (red) detect SEO1-HA which is confined to plasma membrane and colocalizes (merged) with plamsa membrane marker anti-PMA1 antibody (green).

We further estimated the degradation of γ-Glu-Met into glutamate and methionine (i.e. the ability of cells to metabolize γ-Glu-Met). SEO1 was expressed from a TEF promoter in *dug3Δ met15Δ* and *ecm38Δ met15Δ* strains separately, and the accumulation of γ-Glu-Met was monitored using LC-MS/MS. These two strains are defective in γ-Glu-Met degradation (based on our growth experiments). We observe that these strains had significantly lower methionine levels, establishing the requirement of DUG3/DUG2 complex and ECM38 in γ-Glu-Met degradation. Glutamate levels were also reduced in *ecm38Δ and dug3Δ* cells in comparison to *met15Δ*, however this was not as significant as the decrease in methionine levels (Fig 3c). This is expected, since substantial amounts of glutamate will be synthesized *de novo* by cells.

The physiological and uptake experiments suggested that Seo1p must be present at the plasma membrane. To confirm this and to assess any other intracellular localization, we tagged SEO1 with c-terminal HA-tag. The tagged clone was functional (Fig S2) and the localization studies indicate that the protein was predominantly localized to the plasma membrane (as observed with colocalization with the PMA1, a plasma membrane ATPase) (Fig 3d). Seo1p is a member of the Dal5p family of permeases. This family protein members have 12 transmembrane domains, and includes the high affinity cysteine transporter Yct1p, Dal5p etc. SEO1 was first identified in a screen for mutants resistant to the methionine analogue, ethionine sulfoxide, but the substrate for this transporter has remained elusive till now.

### Evaluation of different analogues and related compounds reveals Seo1p is a specific transporter of γ-Glu-met

We next asked if other analogues of γ-Glu-Met could also be transported by Seo1 and examined di-peptides that are structurally similar to γ-Glu-met. Since γ-Glu-cys, is an intermediate of the glutathione cycle and found to be present in yeast cells, we first examined how this peptide is utilized. We observed that it could be utilized as a sulfur source, but in this case, its transport was dependant on the glutathione transporter OPT1/HGT1 and not on SEO1 (Fig 4b). Other γ- Glu peptides such as γ-Glu-leu and γ-Glu-his were also evaluated. To study the growth on γ- Glu-leu and γ-Glu-his, leucine and histidine auxotrophic strains were used for respective peptides. However, the strains were unable to grow on these peptides. The reason for this could be either the absence of a transporter for γ-Glu-leu and γ-Glu-his or the absence of a degradation pathway to provide leucine and histidine for cell growth. Therefore, a competitive uptake inhibition of SEO1 was carried out using an excess of γ-Glu-leu (400uM) along with γ- Glu-met (50uM), and γ-Glu-Met accumulation was estimated using LC-MS. However, cells did not show any reduction in γ-Glu-Met, uptake in the presence of γ-Glu-Leu suggesting that SEO1 is specific to γ-Glu-Met uptake (Fig 4c).

**Figure 4:**
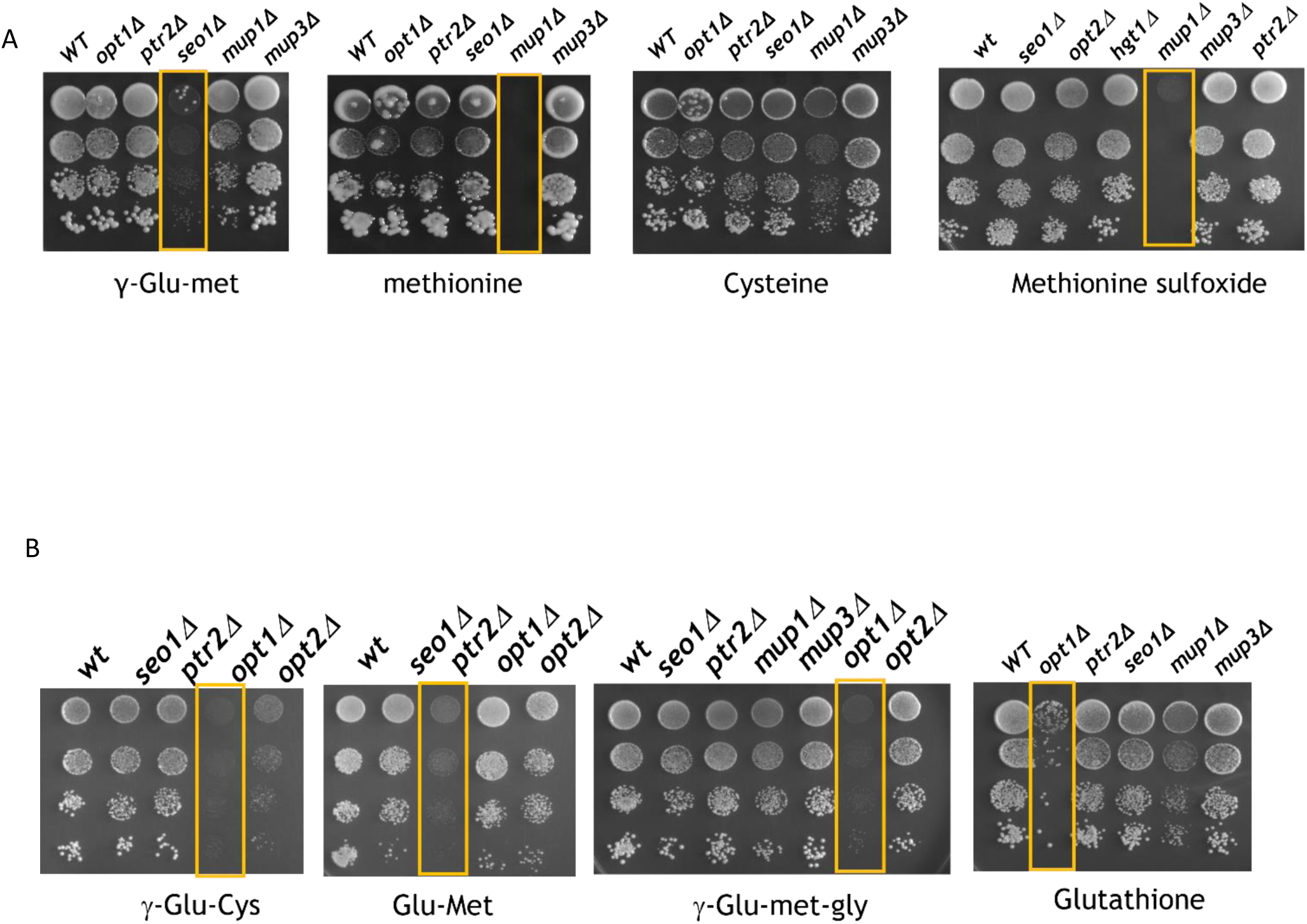
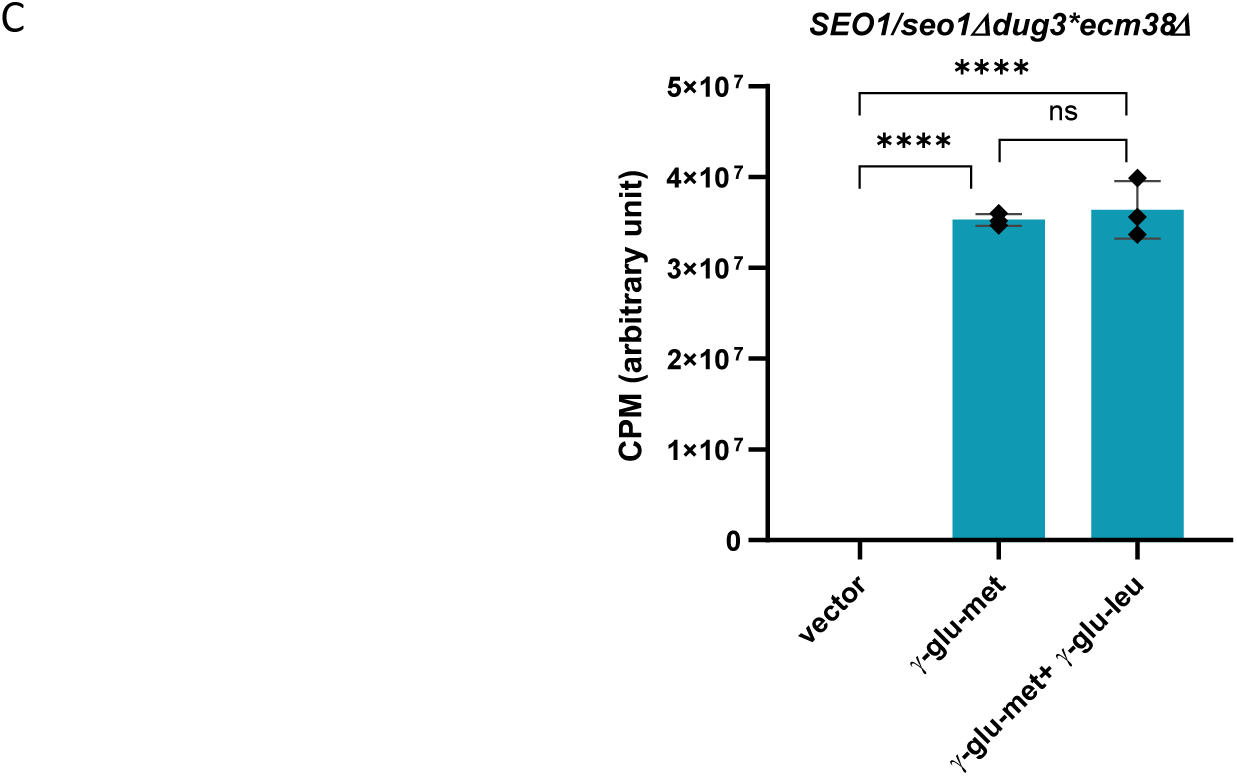
SEO1 transports γ-Glu-met but no other analogous compounds: A) Comparison of different disruptants for utilization of either γ-Glu-met, methionine, cysteine, or methionine sulfoxide. B) Comparison of different disruptants for utilization of either γ-Glu-cys, Glumet, γ-Glu-met-gly or glutathione. **C)** γ-Glu-leu (even at 400μM excess levels) was unable to inhibit uptake of 50 μM γ-Glu-Met by SEO1 in a *seo1Δmet15Δecm38Δdug3-2* background

We also evaluated the tripeptide analogue, γ-Glu-met-gly using growth experiments. We observed that this peptide could support the growth of *seo1Δ* cells well. The only deletion strain that was defective for growth of this tripeptide was the *hgt1Δ /opt1Δ* transporter that is otherwise essential for glutathione transport. These experiments indicated that the γ-Glu-met-gly tri-peptide was not transported by the Seo1p transporter.

When we examined the peptide Glu-met (that contained the normal peptide bond rather than the γ-glutamyl bond), we observed that this peptide was able to support growth of the *met15Δ* cells well. Further, the *seo1Δ* was not defective in growth on this peptide, and the only deletion strain that was defective in growth on this peptide was the strain deleted for the peptide transporter, *ptr2Δ*.

Since *seo1Δ* mutants were initially isolated as mutants resistant to ethionine sulfoxide, we also evaluated methionine sulfoxide as a possible substrate of Seo1p using growth experiments. As can be seen from the figure, the *seo1Δ* was not defective in transport of the methionine sulfoxide. In fact, only the transporter for methionine, Mup1p seemed to be primarily responsible for its utilization as a sulfur source, since only *mup1Δ* were defective on the growth of methionine sulfoxide as a substrate (Fig 4a). Together these studies along with the uptake studies have led us to conclude that the Seo1p transporter is indeed very specific for the γ-Glu-met peptide.

### *Candida auris* and *Candida albicans* SEO1 can also transport γ-Glu-met

We examined for the presence of orthologs of the *S.cerevisiae* Seo1p in other organisms. Seo1 orthologs appeared to be present in many yeasts and fungi, although we could not detect any orthologs in plants and humans. Putative orthologs that were identified in *Candida auris, Candida albicans* were cloned in pRS416TEF plasmid and expressed in *S. cerevisiae seo1Δ* strain (Fig 5a). Uptake of γ-Glu-met (measured using LC/MS/MS) further confirmed the involvement of SEO1 homologs in γ-Glu-met transport (Fig 5b).

**Figure 5:**
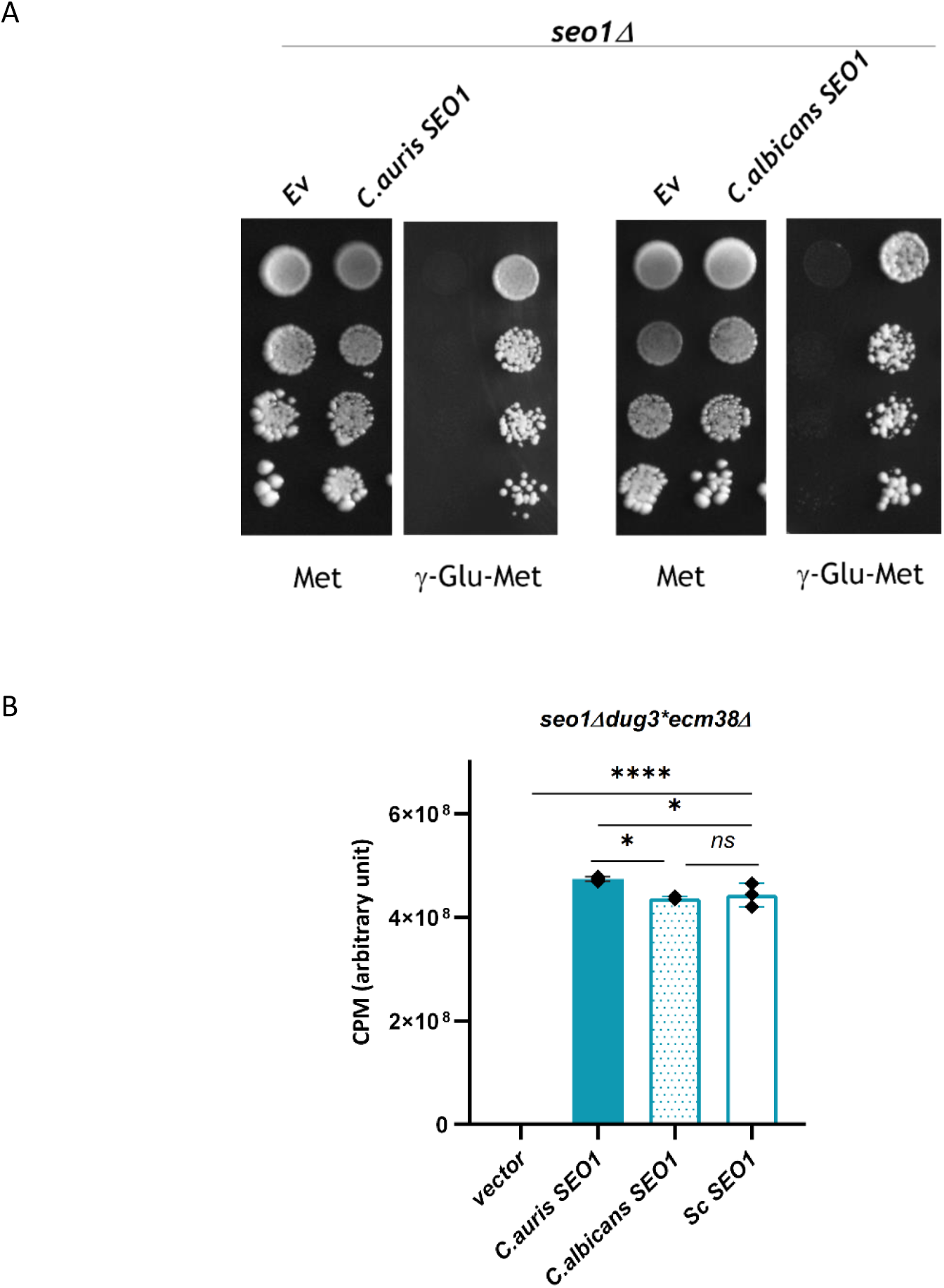
Candida spp SEO1 orthologs encode functional γ-Glu-met transporters: A) *Candida auris* and *Candida albicans* SEO1 expressed from the TEF promoter rescues the defective phenotype of *seo1Δ* for γ-Glu-met, B) confirmation of γ-Glu-met transport by candida SEO1 using LC-MS/MS, these transporters are as efficient as ScSEO1.

### *opt2Δ* affects γ-Glu-met metabolism due to the alteration in vacuolar morphology

In the initial screening of the different transporter deletions for growth on γ-Glu-met, in addition to *seo1Δ* we had observed that deletions in the OPT2 transporter, *opt2Δ,* also showed a fairly severe growth defect (Fig 2b).

As we could clearly establish Seo1p as a plasma membrane transporter for γ-Glu-met, these observations of the *opt2Δ* defect on γ-Glu-met were surprising. OPT2 is a paralog of OPT1/HGT1, a member of the oligopeptide transporter family. OPT1/HGT1 is the yeast glutathione transporter (Bourbouloux et al., 2000). However, the exact function of OPT2 has remained a mystery. An earlier study had localized an N-terminal GFP tagged Opt2p to the punctate organelles that appeared to be peroxisomal and suggested a functions as a peroxisomal and membrane glutathione transporter (Elbaz-Alon et al., 2014). However, other studies including a more detailed investigation revealed that OPT2 is primarily localized to golgi and partly to the plasma membrane and have demonstrated that the *opt2Δ* has a severe vacuolar morphology (Yamauchi et al., 2015). The primary function proposed was in regulating phospholipid asymmetry. To examine if OPT2 might in fact also be a second γ-Glu-met transporter, we examined if OPT2 overexpression could complement the *seo1Δ* phenotype. However, OPT2 failed to complement the *seo1Δ* to any significant extent (Fig 6a). This was further confirmed by the uptake studies (Fig 6b) where we observed only a very small uptake of γ-Glu-met by OPT2 overexpression. This did not explain the strong phenotype of *opt2Δ* on γ-Glu-met. Since Opt2p was suggested to play a role in vacuolar morphology, we examined this phenotype afresh and did indeed observe disrupted vacuolar morphology with numerous small vacuoles in *opt2Δ* (Fig S4). One possible explanation for the severe phenotype of *opt2Δ* on γ-Glu-met therefore was not due to any significant role in transport but as a consequence of defective vacuolar morphology. Ecm38p, γ-Glutamyl transpeptidase, which is involved in γ- Glu-met degradation is a vacuolar enzyme. It is possible that the function of Ecm38p was disrupted in an *opt2Δ* background. To test this hypothesis, we decided to overexpress ECM38 with a strong promoter. We observed that the overexpression of ECM38 in *opt2Δ* strain rescues the defective phenotype on γ-Glu-met. These observations demonstrate that OPT2 has only a minor role in γ-Glu-met transport, and was primarily involved in the functioning of the degradation pathway. We also examined if the overexpression of ECM38 was able to suppress the disrupted vacuolar morphology of *opt2Δ* cells. However, in *opt2Δ* cells bearing the pRSTEF416-ECM38 plasmid we still observed that the cells contained a disrupted vacuolar morphology. This suggested that Ecm38p could only rescue the growth defect of *opt2Δ* on γ- Glu-met and not the other defects. To confirm if the suppression in the growth defect was indeed due to the enzymatic activity of Ecm38p, we used an active site mutant of Ecm38p (G494 → D494) (Kumar et al., 2003) (Fig 6c). However, expressing these active site mutants downstream of the TEF promoter did not lead to growth of *opt2Δ* on γ-Glu-met. This confirmed that increasing the Ecm38p enzymatic activity was responsible for suppressing the *opt2Δ* phenotype on γ-Glu-met.

**Figure 6:**
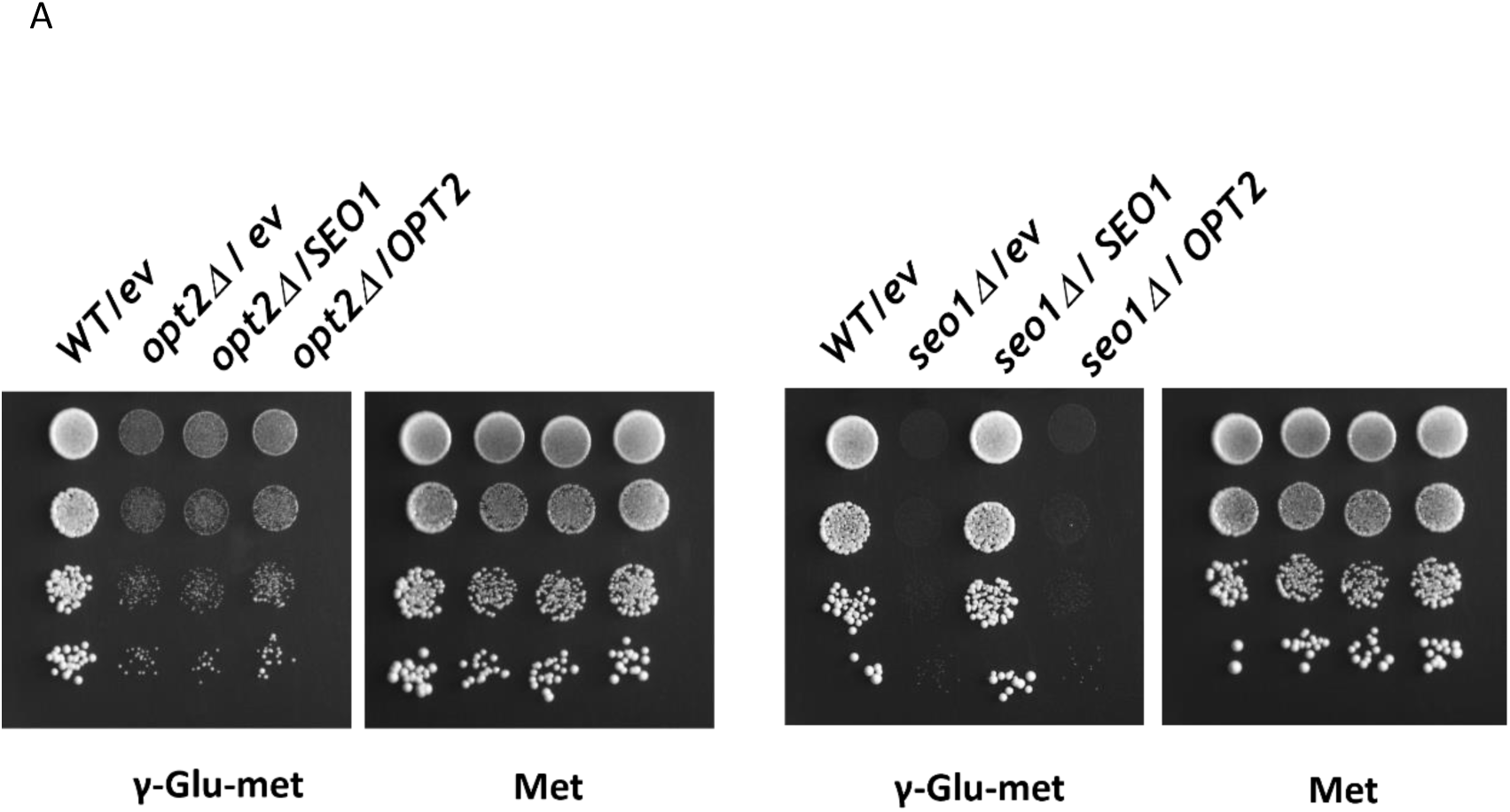
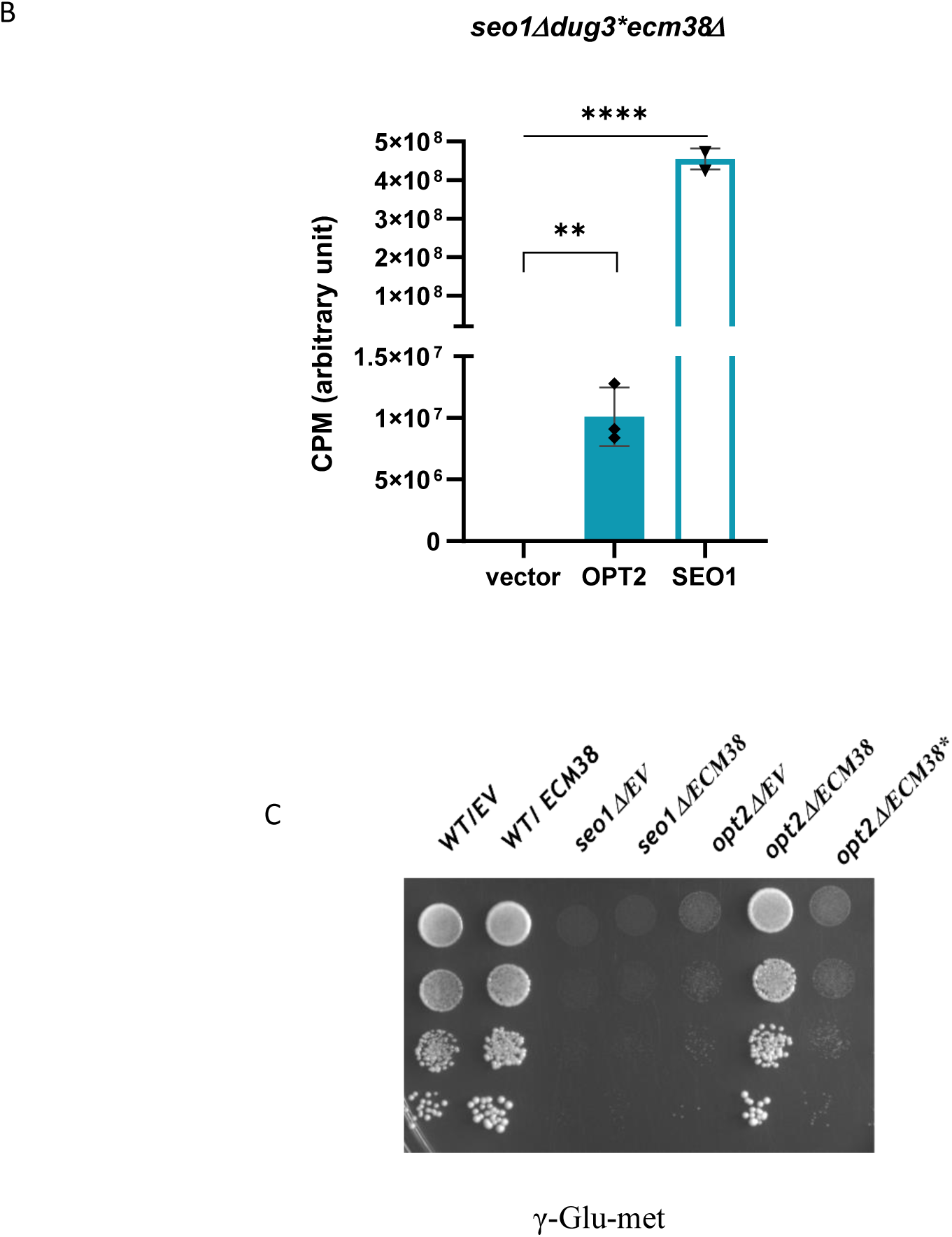
*opt2Δ* interferes with γ-Glu-met utilization not by any role in uptake but by its interference with Ecm38p activity through disruption of vacuolar morphology A) OPT2 overexpression in *seo1Δ* does not rescue the defective phenotype of *seo1Δ* (right panel) whereas there was no rescue seen in *opt2Δ* phenotype upon overexpression of either SEO1 or OPT2 (left panel). B) Weak uptake of γ-Glu-met by OPT2 relative to SEO1. C) ECM38 expressed under TEF promoter rescues the defective phenotype of *opt2Δ* whereas an active site mutant of ECM38 (ECM38*) was unable to rescue the defective phenotype of *opt2Δ* on γ-Glu-met.

### SEO1 is a sulfur regulated protein

In the transcriptomics data we had observed that in growth on γ-Glu-met versus methionine Seo1 was upregulated along with many other sulfur regulated genes. SEO1 has been observed in other genome wide transcription profiles to be upregulated under sulfur limitation. To evaluate this, we first carried out qPCR studies. We observed that the Seo1 mRNA increased in low sulfur conditions (-met) whereas in the presence of methionine the mRNA folds were reduced which suggests that SEO1 expression is sulfur regulated (Fig 7a). We also cloned SEO1 promoter (700bp) upstream of lacZ (β-galactosidase) gene in pLG699z vector to examine β-gal reporter activity in different sulfur conditions. SEO1 was induced under conditions where no methionine was added, whereas in the presence of methionine it was repressed. Similarly, SEO1 was also repressed when cysteine, GSH and even the normal dipeptide n-Glu-met was provided as a sulfur source. In contast, SEO1 was induced in the presence of γ-Glu-met (Fig 7b). This regulation by sulfur was further confirmed at the uptake level by transport experiments, where we observed that cells grown in the presence of methionine (200 μM) led to a lack of ability to take up γ-Glu-met, while growth in low methionine (2μM) allowed the uptake of γ-Glu-met (Fig 7c). This data confirms that the SEO1 is a sulfur regulated protein.

**Figure 7:**
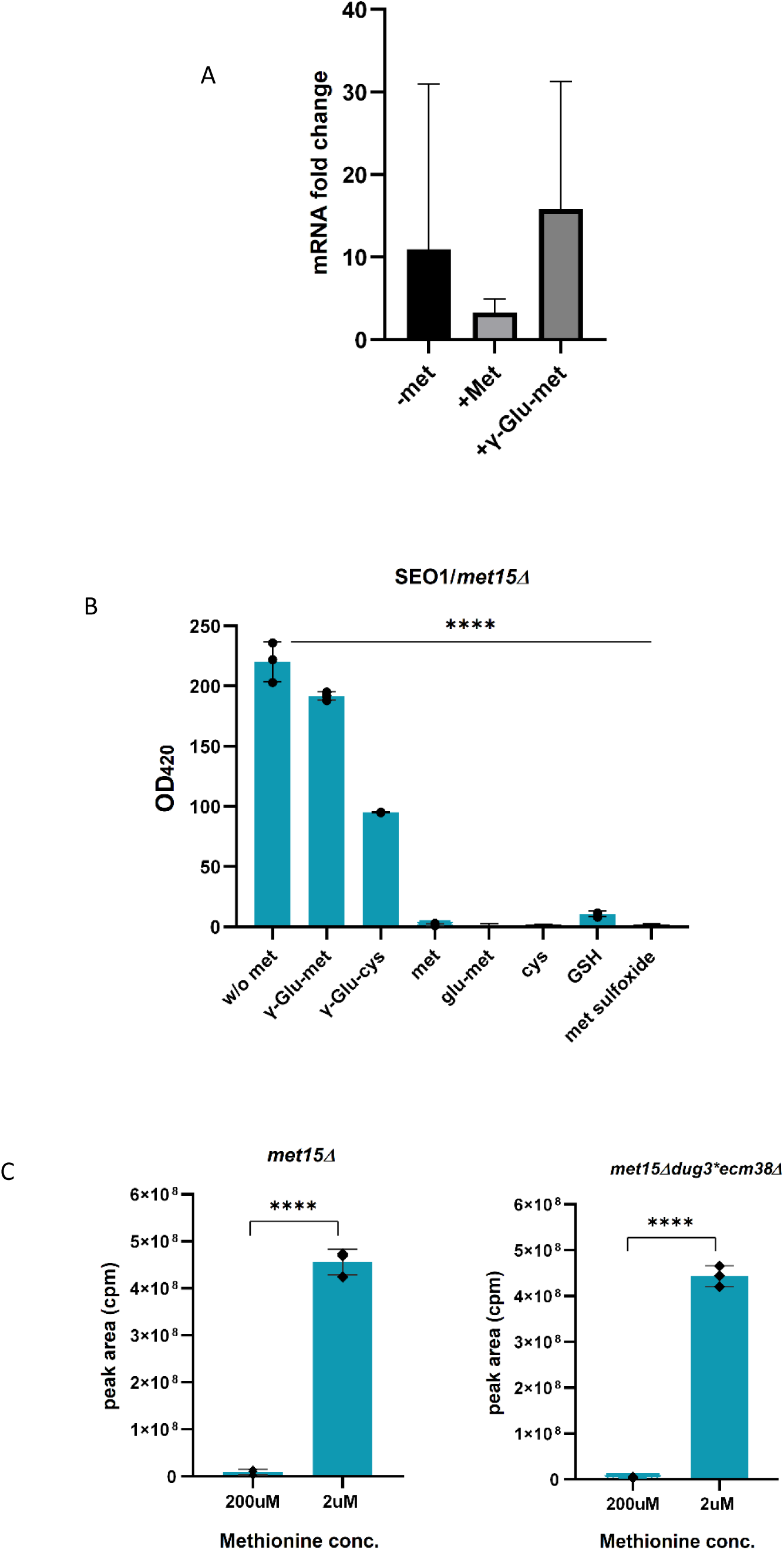
SEO1 is a sulfur regulated protein: A) SEO1 mRNA expression analysis using q-RT-PCR. B) β-galactosidase assay: SEO1 promoter sequence (600bp)-lacZ overexpressed in *met15Δ* strains and it was found to be suppressed in presence of 200μM of methionine, cysteine, GSH, methionine sulfoxide and glu-met. It was derepressed in without methionine or sulfur source condition, 200μM of γ-Glu-met and γ-Glu-cys. C) γ-Glu-met uptake was monitored by LC-MS/MS in presence of high and low concentrations of methionine. 200μM of methionine represses the γ-Glu-met whereas 2μM of methionine does not show the repressing effect.

## Discussion

γ-Glu-di peptides occur ubiquitously; they can thus be used as a nutrient source by different organisms, and can also serve as important metabolites to activate or inhibit pathways. In this study we find that γ-Glu-met can be utilized by yeast as a sulfur source. The growth on γ-Glu-met was only a little slower than on methionine suggesting that it is used as an efficient sulfur source. Investigating its degradation revealed that it is metabolized by two different enzymes; the glutathione degrading enzyme Dug2p/Dug3p complex and the glutathione degrading enzyme Ecm38p. Interestingly, one of them is known to be cytosolic (Dug2p-Dug3p), acting on cytosolic pools, while the other is vacuolar (Ecm38p), acting on vacuolar pools.

What was most surprising in this study, however, was the observation that γ-Glu-met uptake occurs through a dedicated transporter, Seo1p, a previously uncharacterized transporter of the DAL5 family conserved amongst most yeasts and fungi. The high specificity and high affinity (Km for γ-Glu-met was in the micromolar range) was unusual because there are no reports yet of γ-Glu-met being detected in yeasts. A significant effort went into establishing the specificity of Seo1p for γ-Glu-met by testing out other possible substrates. While none of these were substrates of Seo1p, we found out which transporters were involved in the transport of these other compounds. Thus, Mup1p which is known as a high affinity methionine transporter, was also involved in the transport of methionine sulfoxide. The general di/tri peptide transporter Ptr2p was also found to be involved in transport of n-Glu-met. Further, the other tri peptide included in this study, γ-Glu-met-gly, was found to be transported by the very well-known high affinity glutathione transporter Hgt1p.

SEO1 was first isolated in a screen for mutants resistant to ethionine sulfoxide toxicity where MUP1 and MUP3 were also isolated (Isnard et al., 1996). The conclusion the authors could make at that time was that Seo1p does not transport methionine and that it is possibly sulfur regulated. Its isolation as an ethionine sulfoxide mutant further suggested a substrate similar in size or structure. All of these are consistent with γ-Glu-met as the substrate for this transporter. Thus, SEO1 which has for long remained a gene coding for an orphan transporter, has now finally a substrate and function assigned to it.

In this context, it is interesting to note that in a recent study, MFSD1, a lysosomal membrane protein was also recently deorphanized and shown to be a transporter of dipeptides (Jungnickel et al., 2024). Although a variety of dipeptides were evaluated in this study, γ-glutamyl dipeptides were not examined. Since Glu-lys and His-Glu were two of the most efficient substrates transported by MFSD1, given the findings reported here, it would be interesting to see if γ-glutamyl peptides could also be transported.

SEO1 expression was derepressed in the absence of sulfur sources cysteine, methionine, methionine sulfoxide and even n-Glu-met. Strikingly, γ-Glu-met was not a repressing source, while n-Glu-met was a strong repressor of SEO1. This is very consistent with the observation of Seo1p as a γ-Glu-met transporter. Importantly, the ability to transport γ-Glu-met by Seo1p is well conserved across budding yeast, and is not limited to *S. cerevisiae*, since the orthologs from the distant pathogenic yeasts, *Candida auris* and *Candida albicans* were functional in γ- Glu-met transport. Thus, SEO1 seems to be an evolutionarily conserved transporter in the yeasts and fungi.

We also evaluated the role of the putative oligopeptide transporter Opt2p, a homolog of glutathione transporter (Hgt1p/Opt1p) in the utilization of γ-Glu-met, and the deletions of OPT2 showed a defect on the growth of γ-Glu-met. We initially looked at the transport capabilities of this protein. However, uptake data clearly indicated that Opt2p did not import γ-Glu-met significantly. Since the defect of *opt2Δ* for growth on γ-Glu-met was severe, we sought for an alternative explanation and probed the possibility that the reported role of OPT2 in vacuolar function could be important. According to previous reports, OPT2 has a role in phosopholipid asymmetry and affects the assembly of vacuole and maintenance of vacuolar morphology (Yamauchi et al., 2015). We examined if disrupting vacuolar function might be in respect to the γ-Glu-met degrading enzyme Ecm38p which localizes to vacuolar membrane, with its active site facing the vacuolar lumen. While overexpressing this enzyme in an *opt2Δ* background could suppress the growth defect on γ-Glu-met, the disrupted vacuolar phenotype could not be suppressed by this overexpression, and could explain the effects of opt2*Δ* as being due to a deficiency of the Ecm38p activity.

Why yeasts may have evolved a high affinity γ-Glu-met transporter is puzzling. γ-Glu-met is present in human and mouse (Yang et al., 2018a), but there is no report yet of its presence in yeast. A possible hypothesis for the presence of a γ-Glu-met transporter in yeast may lie in the fact that γ-Glu-met is a sulfur containing di-peptide. Sulfur-containing γ-Glutamyl peptides derivatives are widely found in plants (as a defense mechanism against virus, bacterial, microorganisms, insects, predators, and other plants (Amino et al., 2018; Iciek et al., 2009). It is entirely possible that some fungi have evolved a transport mechanism for sulfur containing γ-Glu-di-peptide to salvage whatever sulfur there is in the environment, for sulfur is always limiting. Thus, a sulfur-regulated transporter for γ-Glu-met could be very beneficial in sulfur limiting environments.

## Materials and methods

### Chemicals and Reagents

All chemicals used were of analytical reagent grade. Media components were purchased from Hi Media (India), Merck (Germany) and BD Difco (USA). Oligonucleotides were purchased from Merck (Germany) and GCC (India). Restriction enzymes, Phusion polymerase, dNTPs and other modifying enzymes were obtained from New England Biolabs (Beverly, MA, USA). Gel-extraction kits and plasmid miniprep columns were obtained from QIAGEN (Germany) or Thermo-Fischer Scientific (USA). For initial experiments γ-Glu-met was custom synthesized from Bachem (Switzerland) and for further experiments γ-Glu-met and other γ-Glu-di-/tri-peptides were custom synthesized from GenScript (Biotech Desk Pvt. Ltd. Hyderabad, India). Antibodies were purchased from AbCam (UK) and Cell signalling technologies (USA).

### Strains, media and Growth

The Escherichia coli strain DH5a was used as a cloning host. The list of yeast strains used in this study is shown in Table S3. Yeasts were routinely maintained on YPD medium. The minimal medium contained YNB, glucose, and ammonium sulfate supplemented with the required amino acids and bases. Yeast DNA Isolation and Yeast Transformation—Yeast chromosomal DNA was isolated by the glass bead lysis method and yeast transformations were carried out using the lithium acetate method (Gietz and Woods, 2002).

## Growth curve in liquid culture

The sulfur auxotrophic strain *(met15Δ)* was grown in complete synthetic medium overnight. Further, the inoculum (25 ml) was initiated at 0.01 OD_600_ on complete synthetic medium supplemented with 100μM γ-Glu-met, 100μM methionine (control) as sulfur source and growth was monitored for 32 hours. The data were plotted to obtain the generation time during logarithmic growth.

## RNA sequencing

The AB6302 strain was grown in SD medium containing 100μM methionine (control) and 100μM γ-Glu-met (test) as a sulfur source. The mid log phase cells were harvested, washed with water and frozen. Total RNA was isolated using Qiagen RNeasy mini kit (Cat # 74106), RNA sequencing libraries were prepared with Illumina-compatible NEBNext® Ultra™ II Directional RNA Library Prep Kit (New England BioLabs, MA, USA) were carried out by Genotypic Pvt.Ltd.. Further, paired-end Illumina Next Generation Sequencing was performed at Genotypic Technology Pvt. Ltd., Bangalore, India.

## Recombinant DNA

SEO1 (1700bp) gene was cloned in pRS416TEF by PCR amplification followed by homologous recombination in yeast as described previously (Singh et al., 2024). The gene specific primers carried overlapping sequences for pRS416TEF and an internal restriction site Hind III and XhoI were also added in forward and reverse primers respectively (Table S2). Two further sets of primers were also designed to amplify the vector in two different fragments carrying the overlapping sequences for homologous recombination. One fragment of the vector called CEN fragment as it contains CEN another called URA fragment as it contains URA marker. For homologous recombination the CEN and URA fragments along with the SEO1 gene fragment, carrying vector overlapping sequences, were transformed in yeast using lithium acetate method and plated on SD-URA plates. A few colonies were picked up and desired recombinants were confirmed by PCR. The Plasmids were isolated after passage through E. coli and digestion by the restriction enzymes HindIII and XhoI confirmed the presence of the clone. OPT2, OPT2-HA and Candida SEO1 were also cloned using the same homologous recombination method and the primers used are described in Table S2. *C. auris* and *C. albicans* have a slightly different genetic code in the LEU codon. The SEO1 gene of both these yeasts had a single codon which codes for leucine in *Candida albicans* and *Candida auris* but serine in *Saccharomyces cerevisiae*, but were cloned and expressed without correcting for the codon.

## Growth assay by dilution spotting

For growth assay, the different strains were grown overnight in minimal ammonia medium without uracil and reinoculated in fresh medium to an OD600 of 0.1 without Uracil and methionine and grown for 6 hr. The exponential-phase cells were harvested, washed with water, and resuspended in water to an OD600 of 0.2. These were serially diluted to 1:10, 1:100, and 1:1000. Of these cell resuspensions, 10 ml were spotted on minimal medium different sulfur sources described as sole organic sulfur source. The plates were incubated at 30 degrees for 3 days and photographs were taken.

## Cellular localization of SEO1 using microscopy

c-terminal HA tagged pRS416TEF-SEO1 was transformed in *seo1Δ* strain, these transformants were further growth in SD without uracil medium for localization experiments. Primary culture was grown overnight and further secondary culture was started at 0.2OD and grown for 4-5 hrs. The cells in exponential phase were harvested and spheroplasts were made using zymolyase. The spheroplasts were washed with KPO4 buffer and fixed in 4% PFA. Further, permeabilization was done by using 0.1% Triton X and blocked with 1% BSA. Cells were further incubated with Anti- rabbit Anti-HA antibodies (1:1000 dilution) and Anti-mouse Anti- PMA1 (1:1000 dilution) antibodies at 4 degrees overnight. Next day, cells were washed with PBS and incubated with Anti-mouse Alexa flour 488 (1:500 dilution) and Anti-rabbit Alexa flour 647 (1:500 dilution) secondary antibodies for 2 hrs at RT. Finally, cells were washed with PBS and visualized under Nikon ti2 eclipse microscope.

## Vacuolar morphology and microscopy

To visualize the vacuole, cells in log phase were stained with 8μM FM4-64 (Cat # S6689) for 30 min, washed and resuspended in the SD medium, and incubated for an additional 1 hour at 30°C (Yamauchi et al., 2015). cells were visualized under Nikon ti2 eclipse microscope.

## Metabolite extraction and LC-MS/MS analysis

To measure the uptake of γ-Glu-met peptide by SEO1, *seo1Δ dug3-2 ecm38Δ* (AB6601) cells with SEO1 overexpression and empty vector (control) were grown in SD without uracil medium overnight and reinoculated in SD without uracil and methionine medium at 0.02 OD_600_ for 4-5 hours. Cells in exponential phase were supplemented with 100μm γ-Glu-met, metabolite quenching and cell harvesting were done at different time intervals i.e. 5, 10, 20, 40 and 60mins. The cells were pelleted down at 600g for 3 min, and metabolites were extracted, as described earlier (Walvekar et al., 2018). Briefly, 1 ml of ice-cold 10% methanol was added without disturbing the pellet (to quench metabolism) and further centrifuged at 600g for 3 min at 4°C. Furthermore, 1 ml of 80% methanol (maintained at −45°C) was added to the pellet, vortexed for 15 s, and incubated at −45°C for 15 min for metabolite extraction. The tubes were vortexed (15 s) and centrifuged at 21,000g for 10 min at −5°C. The supernatant (900 μl) was transferred into fresh tubes, recentrifuged at 21,000g for 10 min at −5°C, and the supernatant was removed and dried using a SpeedVac. The samples were stored at −80°C briefly, before analysis by targeted LC/MS/MS to assess the γ-Glu-met uptake, expanding methods described earlier (Walvekar et al., 2018). We used Synergi TM 4-m Fusion-RP 80-Å (100, 4.6 mm) LC column for detection, and the indicated metabolites were estimated using an ABSciex QTRAP 6500 in triple-quadrupole mode. For kinetics experiments 10, 50, 100, 150 and 200μM γ-Glu-met was used for uptake and cells were harvested at 40 min. A similar protocol was followed for uptake of γ-Glu-met by candida SEO1 and ScOPT2 in *seo1Δ dug3-2 ecm38Δ* strain at 40 mins time point.

## q-RT-PCR to study SEO1 regulation

Yeast cells were grown with -met, +met and +γ-Glu-Met supplementation and harvested in exponential phase of growth. RNA was isolated from these cells using Acid-Phenol method and cDNA was prepared using Thermo Fischer Scientific RevertAid cDNA synthesis kit. For genes, the primer set listed in Table S2 were used with a final reaction volume of 5μl containing using Maxima SYBR Green qPCR Master Mix (Fermentas, USA). PCR conditions were 95°C for 3 min, then 40 cycles consisting of denaturation at 95°C for 10 s, annealing at 60°C for 10s and extension at 72°C for 30 s, followed by the melting curve protocol with 10s at 95°C and then 60s each at 0.5°C increments between 65°C and 95°C. The reactions were performed in triplicate for each sample. The relative amounts of target gene expression for each sample were calculated using the Livak method formula 2-(ΔΔCT) against an endogenous control actin for genes. Finally, the fold change against the control gene is calculated and plotted. Data analysis was performed by Graph pad prism using one way ANOVA. The significance of differences between means were calculated at a 5% level (P< 0.05).

## Construction of the SEO1 promoter–LacZ fusion constructs for the β-galactosidase reporter assay

The SEO1 promoter sequence (600bp) was PCR amplified using oligonucleotides listed in Table S2. The PCR products were purified, digested with XhoI and BamHI, and cloned into pLG669z (Guarente and Ptashne, 1981). pLG669z -SEO1-Promoter was transformed into the *met15Δ* (AB5000) strain and the transformants were selected on minimal media plates without uracil. Further, these transformants were grown in presence of different sulfur sources such as 200μM of methionine, γ-Glu-met, GSH, methionine sulfoxide, cysteine, glu-met and γ-Glu-cys, as indicated. The β-galactosidase reporter assay was carried out using standard protocols, and relative activity presented as OD420.

## Induction conditions and β-galactosidase assay

Fresh yeast transformants were used in all β-galactosidase experiments. The transformants were picked, grown overnight in minimal ammonia medium without uracil, and reinoculated in fresh minimal ammonia medium (without any organic sulfur source) to an initial OD600 of 0.2. They were grown for an additional 6–7 hr to induce β-galactosidase. The cells were then harvested and washed twice in LacZ buffer (60 mm Na2HPO47H2O, 40 mm NaH2PO4H2O, 10 mm KCl, 1 mm MgSO47H2O, 0.27% 2-mercaptoethanol, pH 7) and taken for the reporter gene assay. β-Galactosidase activity was assayed in permeabilized yeast cells essentially as described previously (Guarente and Ptashne, 1981). Briefly, the cells were resuspended in LacZ buffer, permeabilized by the addition of 50 ml chloroform and 20 ml SDS (0.1%). They were then vigorously vortexed for 20 sec. These samples were equilibrated for 10 min at 30 and then o-nitrophenyl β-galactopyranoside was added sequentially to the reaction samples. β- Galactosidase units are given as OD420 × 1000 min1 ml1 / OD600 at 30. The experiments were repeated with a minimum of three independent colonies.

## Statistical analysis

All uptake data has been analysed using student’s t test and one-way Anova with p value cut off of 0.05 using Graph Pad Prisma. Standard kinetics-based Km determination was applied.

## Author contributions

PD: Performed most of the experiments, data analysis and manuscript writing; MS: performed LC-MS/MS experiments; SL: supervised the design and analysis of the uptake experiments, and part of the manuscript writing; AKB: Supervision, experiment designing, conceptualization and manuscript writing.

## Acknowledgement

The work was supported by a grant-in-aid project from the Department of Biotechnology (BT/PR39040/BRB/10/1882/2020) to AKB and a DBT-Wellcome Trust India Alliance Senior Fellowship (IA/S/21/2/505922) to SL. PD was a recipient of a Senior Research Fellowship from the Indian Council of Medical Research (ICMR), MS thanks to SERB NPDF2022(PDF/2022/000700) for funding. We thank NCBS/InStem/CCAMP Mass Spectrometry facility for liquid chromatography-tandem mass spectrometry used for the uptake studies. We thank Shravan Kumar Mishra and his Lab for help with the microscope facility. We thank Ganesh Muthu for help with the RNAseq analysis. We also thank Dr Praveen Singh and Dr Shantanu Sengupta, Institute of Genomics and Integrative Biology, New Delhi for help with some of the initial mass spec analysis.

## Author affiliations

^1^Indian Institute of Science Education and Research, Mohali, Punjab, India, ^2^ DBT- Institute for Stem Cell Science and Regenerative Medicine (inStem), Bangalore, India,

